# UPL3 Promotes BZR1 Degradation, Growth Arrest, and Seedling Survival under Starvation Stress in *Arabidopsis*

**DOI:** 10.1101/2023.10.18.562997

**Authors:** Zhenzhen Zhang, Hongliang Zhang, Efren Gonzalez, Tarabryn Grismer, Shou-Ling Xu, Zhi-Yong Wang

## Abstract

BRASSINAZONE RESISTANT 1 (BZR1) is a key transcription factor of the brassinosteroid signaling pathway but also a signaling hub that integrates diverse signals that modulate plant growth. Previous studies have shown that starvation causes BZR1 degradation, but the underlying mechanisms are not understood. Here we performed quantitative proteomic analysis of BZR1 interactome under starvation conditions and identified two BZR1-interacting ubiquitin ligases, BAF1 and UPL3. Compared to the wild type, the *upl3* mutants show long hypocotyl and increased BZR1 levels when grown under sugar starvation conditions but not when grown on sugar-containing media, indicating a role of UPL3 in BZR1 degradation specifically under starvation conditions. The *upl3* mutants showed a reduced survival rate after starvation treatment, supporting the importance of UPL3-mediated BZR1 degradation and growth arrest for starvation survival. Treatments with inhibitors of TARGET of RAPAMYCIN (TOR) and autophagy altered BZR1 level in the wild type but were less effective in *upl3*, suggesting that UPL3 mediates the TOR-regulated and autophagy-dependent degradation of BZR1. Further, the UPL3 protein level is increased posttranscriptionally by starvation but decreased by sugar treatment. Our study identifies UPL3 as a key component that mediates sugar regulation of hormone signaling pathways, important for optimal growth and survival in plants.

**IN A NUTSHELL:** *Background:* The coordination between signaling pathways that monitor the levels of photosynthate and growth hormones is crucial for optimizing growth and survival, but the underlying mechanisms are not fully understood. When the sugar level is low, the BZR1 transcription factor of the brassinosteroid (BR) signaling pathway is degraded, and hence growth is attenuated to prevent starvation and enhance survival. When sugar is sufficient, sugar signaling inhibits BZR1 degradation and enables BR promotion of plant growth. The key component that mediates starvation-induced BZR1 degradation remains unknown.

*Question:* What proteins interact with BZR1 and mediate its degradation under sugar starvation?

*Finding:* We performed immunoprecipitation mass spectrometry analysis of BZR1 in starvation-treated Arabidopsis and identified many BZR1-interacting proteins, including two E3 ligases UPL3 and BAF1. Genetic analysis showed that UPL3 plays a specific and prominent role in promoting autophagy-dependent BZR1 degradation and plant survival under sugar-starvation conditions.

*Next step:* How sugar-TOR signaling regulates UPL3 level remains to be studied in the future.

## Introduction

Brassinosteroids (BRs) are a class of growth hormone that regulates development and acclimation to environmental conditions throughout the life cycle of plants (Clouse and Sasse, 1998; Chaiwanon et al., 2016; Nolan et al., 2017a; Ortiz-Morea et al., 2020). BRs are perceived by the plasma membrane-localized receptor kinases BRASSINOSTEROID INSENSITIVE1 (BRI1) (Li and Chory, 1997; Wang et al., 2001) which initiates a signaling cascade that leads to activation of the BRASSINAZOLE-RESISTANT1 (BZR1) family transcription factors including BZR1 and BZR2 (also named BRI1-EMS-SUPPRESSOR1, BES1) (Wang et al., 2002). When the BR level is low, BZR1 and BZR2 are phosphorylated by the GSK3-like kinase BRASSINOSTEROID-INSENSITIVE2 (BIN2) (He et al., 2002; Yin et al., 2002), and consequently excluded from the nucleus and unable to bind DNA (Vert and Chory, 2006; Gampala et al., 2007). When the BR level is high, BR induces BRI1 dimerization with and activation by its co-receptor kinase BRI1-ASSOCIATED KINASE1 (BAK1). Activated BRI1 phosphorylates the BR SIGNALING KINASES (BSKs) and CONSTITUTIVE DIFFERENTIAL GROWTH1 (CDG1) kinases, which activate the BRI1 SUPPRESSOR1 (BSU1) phosphatase (Tang et al., 2008; Kim et al., 2011). BSU1-mediated dephosphorylation of BIN2 inhibits its kinase activity and causes its ubiquitination mediated by the KINK SUPPRESSED in *bzr1-1D* (KIB1) ubiquitin ligase (Zhu et al., 2017). Upon inactivation of BIN2, BZR1 and BZR2/BES1 are dephosphorylated by PROTEIN PHOSPHATASE 2A (PP2A) (Tang et al., 2011), and then accumulate in the nucleus to regulate BR-responsive gene expression (He et al., 2005; Yin et al., 2005).

The BZR1 family transcription factors are essential for BR regulation of gene expression and plant growth and development. Deficiency in BR biosynthesis or sensing causes pleiotropic growth defects including extreme dwarfism, photomorphogenesis in the dark, delayed seed germination and flowering, reduced root growth, altered shoot architecture, and male sterility. The dominant gain-of-function mutation *bzr1-1D* suppresses nearly all the phenotypes of BR deficiency (Wang et al., 2002), whereas the *bzr-h* mutants lacking all six members of the BZR1 family show nearly identical developmental and gene-expression phenotypes as the mutants lacking all BR receptors (BRI1 and its two homologs) (Chen et al., 2019). BZR1 directly activates and represses thousands of genes, which contribute to cell elongation and the development of shoot, root, and floral transition, and reproductive cells (Luo et al., 2010; Sun et al., 2010; Yu et al., 2011; Gendron et al., 2012; Li et al., 2018; Cai et al., 2023).

BZR1 homologs appear to play similar growth-promoting roles in crops. Genetic and transgenic experiments have shown functions of BZR1 homologs in regulating leaf angle (Zhang et al., 2009) and grain length in rice (Du et al., 2023), nodulation in soybean, organ size including kernel size in maize (Zea mays) (Zhang et al., 2020; Sun et al., 2021), and growth and salt tolerance in tomato (Jia et al., 2021). A recent study of genome-wide ZmBZR1 binding revealed extensive genetic and epigenetic variations in ZmBZR1 binding, many of which are associated with major traits in maize (Hartwig et al., 2023).

BZR1 is a central hub that integrates diverse internal and external cues in regulating growth, development, and environmental acclimation. In addition to regulation by BR-dependent phosphorylation, BZR1 activity is modulated by other signaling pathways and as such BZR1 mediates growth regulation by other internal and external signals. For example, BZR1 plays an essential role in skotomorphogenesis and cell elongation responses to auxin, gibberellin, and darkness, through interactions with AUXIN RESPONSE FACTOR 6 (ARF6), the gibberellin-signaling component DELLA, and PHYTOCHROME-INTERACTING FACTOR 4 (PIF4) (Bai et al., 2012; Oh et al., 2012; Oh et al., 2014). BZR1 regulates apical hook development and cotyledon greening by interacting with SMALL AUXIN UP RNA17 (SAUR17), PIFs, ETHYLENE INSENSITIVE 3 (EIN3), and EIN3-LIKE 1 (EIL1) and GROWTH REGULATING FACTOR 7 (GRF7) (Li and He, 2016; Wang et al., 2020; Zhao et al., 2021; Wang et al., 2023). BZR1 and BZR2/BES1 mediate hypocotyl elongation responses to UV and blue light by interacting with the photoreceptors UV RESISTANCE 8 (UVR8) and CRYPTOCHROME 1 (CRY1) (Liang et al., 2018; He et al., 2019). BZR1 and BZR2/BES1 modulate biotic and abiotic stress responses by interacting with stress-responsive factors (Lozano-Duran et al., 2013; Chen et al., 2017; Ye et al., 2017; Qi et al., 2021). BZR1 is also regulated by redox-regulated oxidation and it promotes stomatal opening by integrating with the redox signal hydrogen peroxide (Tian et al., 2018; Li et al., 2020).

Degradation of BZR1 and BZR2/BES1 has emerged as a major mechanism of growth inhibition by stresses and sugar starvation. As the energy source, building blocks for cell walls, and product of photosynthesis, sugar is crucial for plant growth and survival. To achieve optimal growth and avoid cellular starvation, growth hormone signaling must be controlled by sugar signaling. Several studies have shown that sugar-hormone crosstalk involves starvation-induced BZR1 degradation, which contributes to growth arrest and alleviation of starvation. Sugar signaling through Target of Rapamycin (TOR) stabilizes BZR1, and when cellular sugar level is low, the inactivation of TOR leads to BZR1 degradation through an autophagy-dependent mechanism (Zhang et al., 2016). In addition, BES1 was shown to be degraded by a selective autophagic pathway mediated by ubiquitin receptor DOMINANT SUPPRESSOR OF KAR2 (DSK2) (Nolan et al., 2017b; Wang et al., 2021). Recently, an F-box E3 ligase, named BES1-ASSOCIATED F-BOX1 (BAF1) was reported as a BES1-interacting protein that mediates sugar starvation-induced selective autophagy of BES1. The *baf1* mutant and SINAT-RNAi plants showed a reduced survival rate after starvation treatment but their growth phenotypes under starvation conditions were not analyzed and they still showed sugar starvation-induced BES1/BZR2 degradation (Wang et al., 2021), suggesting that a mechanism independent of SINAT and BAF1 mediates starvation-induced BZR1/BZR2/BES1 degradation and growth arrest.

To identify proteins that interact with BZR1 under starvation conditions, we performed stable isotope labeling followed by Immunoprecipitation and quantitative mass spectrometry (SILIP-MS) analyses of the BZR1 interactome using seedlings grown under a sugar starvation condition. The experiments detected about 28 known BZR1/BES1 interactors and their family members and identified many new candidates of BZR1-interacting proteins which include two E3 ubiquitin ligases, BAF1 and UPL3. Genetic and biochemical experiments provided evidence for a prominent role of UPL3 in BZR1 degradation and growth arrest under sugar-starvation conditions.

## RESULTS

### Identification of BZR1-interacting Protens using Quantitative Proteomics

To identify proteins associated with BZR1, we performed SILIP-MS as outlined in Figure S1A. To capture sufficient amounts of the proteins that mediate BZR1 degradation during sugar starvation, we performed SILIP-MS using a transgenic line that overexpresses BZR1-YFP (*35S:BZR1-YFP).* We grew the *BZR1-YFP* seedling on sugar-free media containing light nitrogen (^14^N) and the *35S:GFP* negative control on heavy nitrogen (^15^N) under constant light for two weeks and then put the seedling in the dark for 24 hours to cause sugar starvation. The nitrogen isotopes were switched in the replicate experiments. After immunoprecipitation, the samples were quality-checked by immuno-blotting and the beads of ^14^N- and ^15^N-labeled sample and control were mixed (Figure S1A-S1B). The proteins were eluted, separated in an SDS-PAGE gel then in-gel digested by trypsin (Figure S1C), and analyzed in an Orbitrap mass spectrometer. Enrichment by BZR1 was quantified based on the ^14^N/^15^N ratios of the identified peptides (Shrestha et al., 2022). MS analysis identified 2819, 2985, or 2549 proteins in the three replicate experiments. A total of 1598 proteins were identified in all three replicates (Supplementary Dataset 1). These include 28 known BZR1 interactors or their family members, which showed over 20x enrichment in the *BZR1-YFP* relative to the *35S:GFP* control in both forward and reverse labeling repeat experiments. Using 20x enrichment and at least two peptides identified in each replicate as the cut-off, the experiment identified 41 additional putative BZR1-interacting proteins (Supplementary Dataset 1).

### UPL3 Interacts with BZR1

The putative BZR1-interacting proteins identified in our IP-MS experiment include two E3 uniquitin ligases: the BES1-Associate F-box 1 (BAF1) and ubiquitin-protein ligases 3 (UPL3) (Figure S2). BAF1 was recently identified by yeast-two-hybrid screen as an F-box E3 ligase for BES1, and UPL3 is a HECT domain-containing family of ubiquitin-protein ligases, previously reported to play roles in trichome development and stress responses (Patra et al., 2013; Wang et al., 2022). To further confirm BZR1 interaction with UPL3, we performed targeted quantification using parallel reaction monitoring (PRM) MS (Figure 1A-1C, and S3) (Reyes et al., 2022). The UPL3 peptides were enriched over 20-fold compared to *35S:GFP* samples in all three repeat experiments, including two repeats with ^14^N-labeled *BZR1-YFP* (Figure 1A-B and S3) and one with ^15^N-labeled *BZR1-YFP* sample (Figure 1C and S3). In contrast, a control protein ACETYL-COA CARBOXYLASE 1 (ACC1) showed a ratio of around 1 in all three experiments (Figure 1A-1C and S3). In addition, we performed co-immunoprecipitation of BZR1-YFP and UPL3-Myc. UPL3-Myc was expressed alone or co-expressed with BZR1-YFP or 35S:GFP in *Nicotiana benthamian* leaves. After immunoprecipitation with anti-GFP antibody (GFP-trap), immunoblotting showed that UPL3-Myc was immunoprecipitated by BZR1-YFP but not GFP (Figure 1D). Together, these results show that UPL3 interacts with BZR1 *in vivo*.

**Figure 1.**
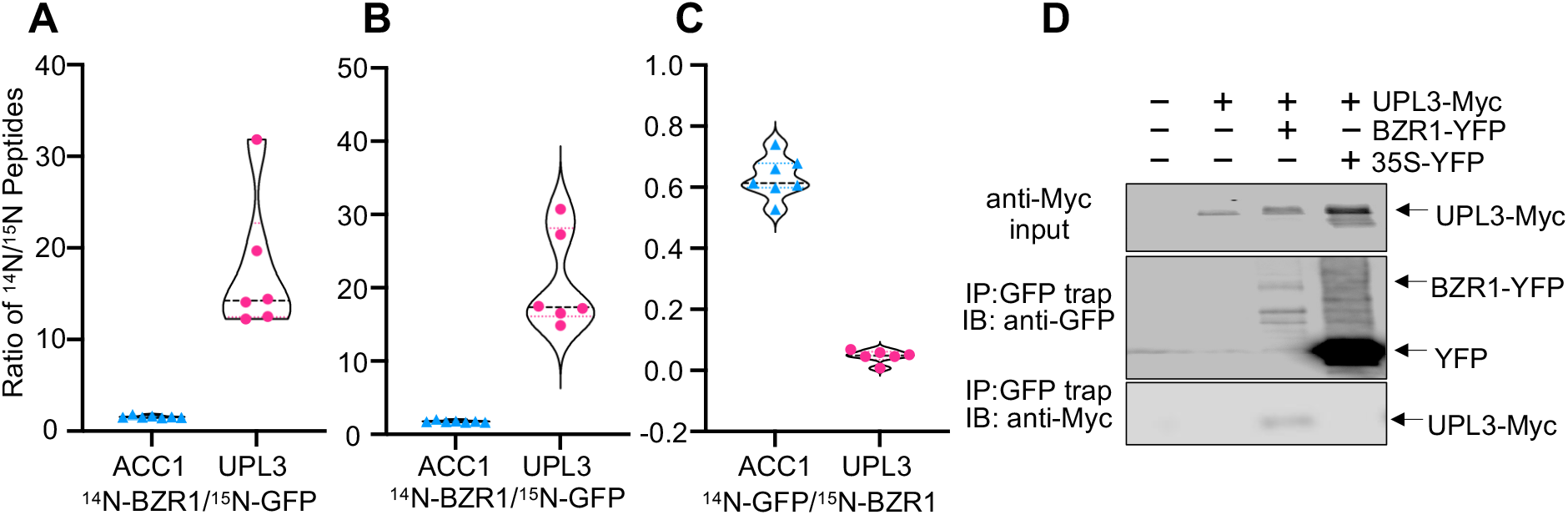
UPL3 interacts with BZR1. **A-C)** PRM quantitation of ^14^N- and ^15^N-labeled peptides of UPL3 and ACC1, as ratios between *BZR1-YFP* and *GFP* control, in three replicates of anti-GFP immunoprecipitation. **D)** Co-IP assay shows *in vivo* interaction between BZR1-YFP and UPL3-Myc. *N. benthamiana* leaves were transiently co-transformed with *BZR1:BZR1-YFP* (or *35S:GFP* as a control) and *35S:UPL3-Myc.* The Input extract and anti-GFP immunoprecipitants (IP) were analyzed by immunoblotting (IB).

**Figure 2.**
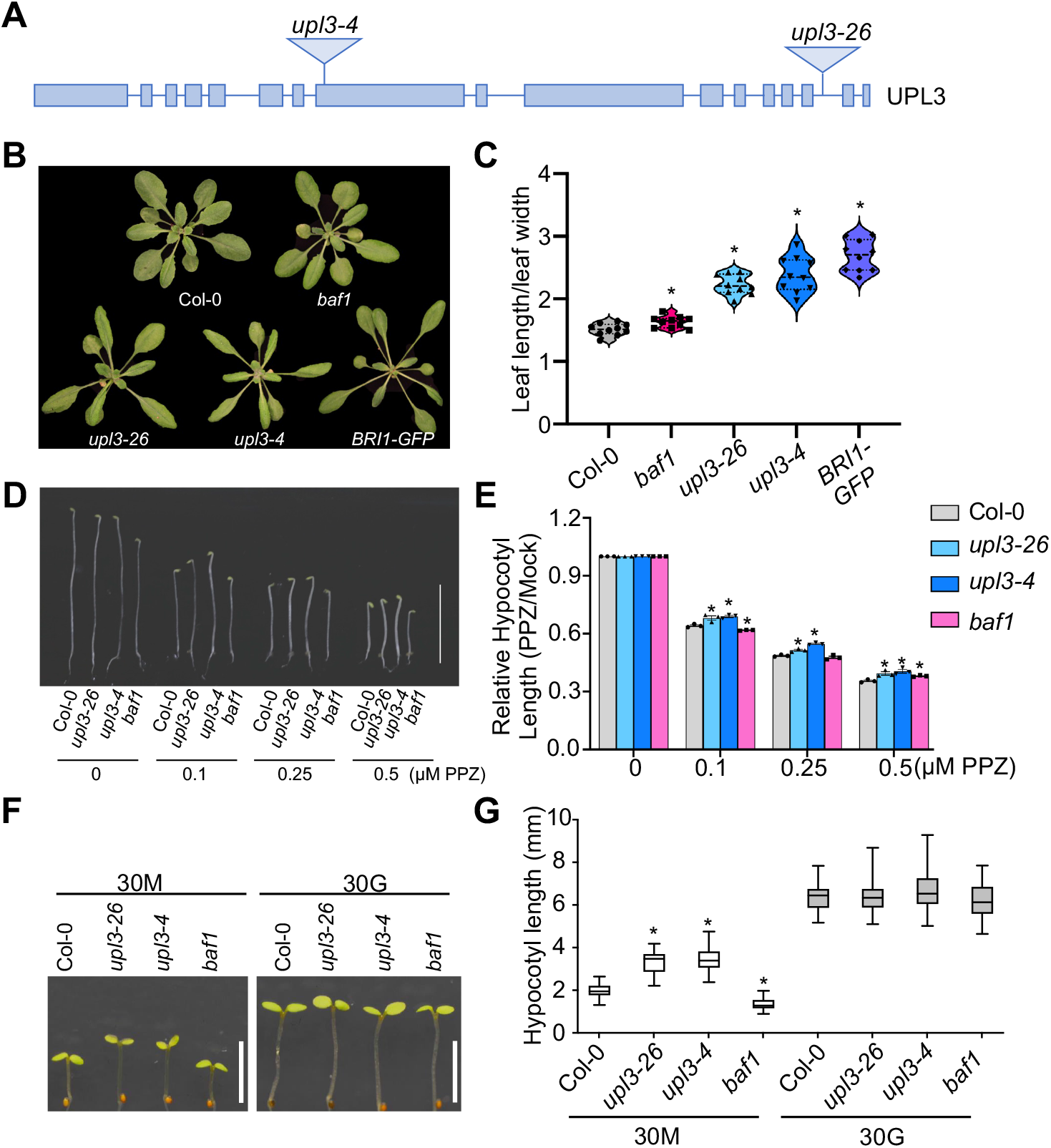
The *upl3* mutants show BR-hyper-response phenotypes. **A)** The T-DNA insertion sites of *upl3-4* and *upl3-26*. **B**) The *upl3-26* and *upl3-4* mutants show elongated petioles and leaves, similar to plants overexpressing BRI1-GFP, when grown in a long-day condition for one month. **C)** Ratios of leaf length to width of the plants represented in panel B. Ten fully expanded leaves of five plants were quantified per genotype, * for *p*<0.05 (Student’s t-test). **D)** The seedlings were grown on media containing the indicated concentrations of PPZ in the dark for 5 days. **E**) The relative hypocotyl length of plants represented in panel D, * for *p*<0.05 (Student’s t-test). **F)** Seedlings were grown under light for 4 days, then in the dark for another 3 days, on media containing 30 mM mannitol (30M) or 30 mM glucose (30G). **G)** Hypocotyl length of seedlings represented in panel F. At least 30 seedlings were included for measurement, * for *p*<0.05 (Student’s t-test).

**Figure 3.**
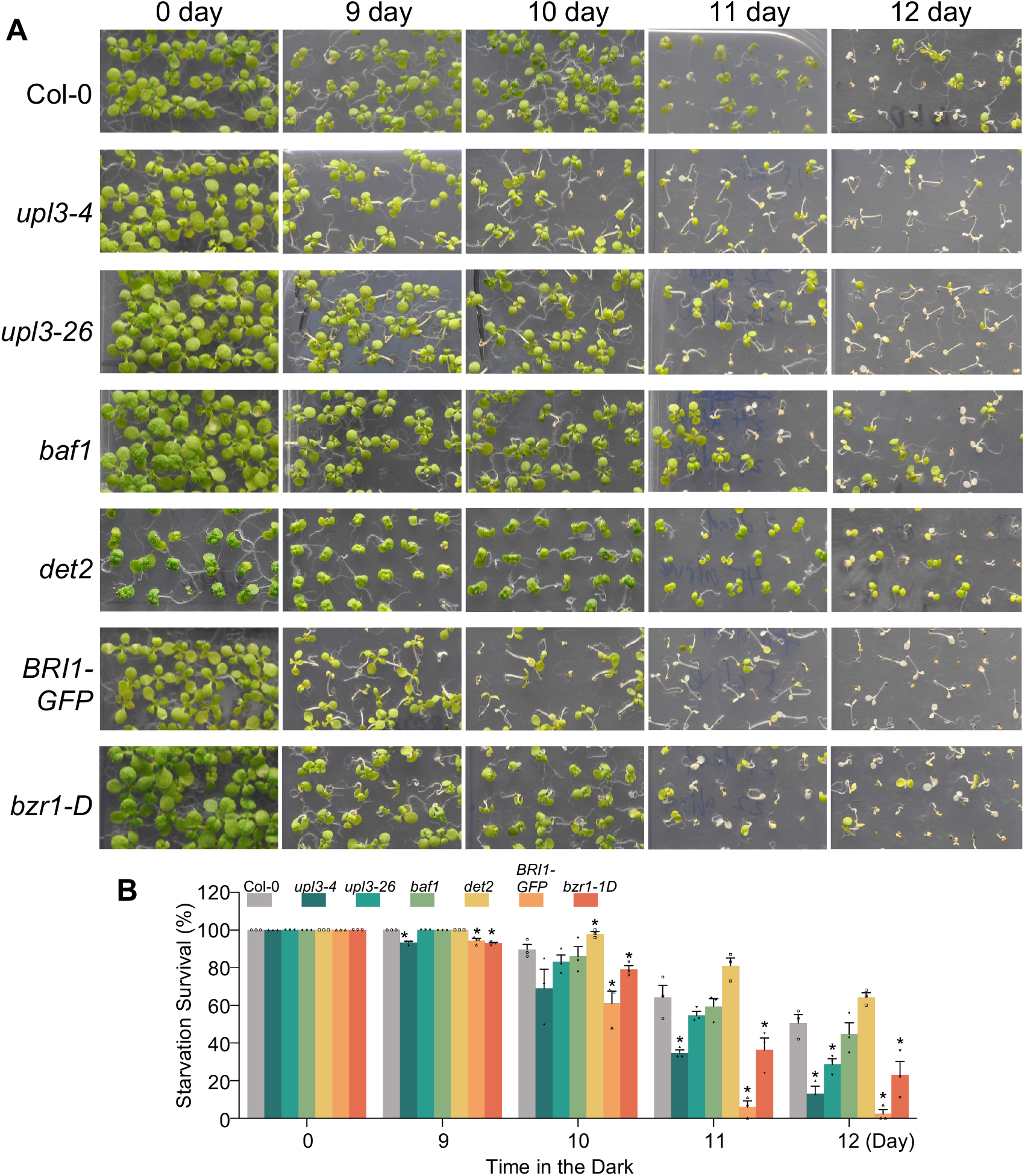
UPL3 promotes starvation survival. **A)** Seedlings were grown on 1/2 MS medium without sugar for 4 days, transferred to the dark for the indicated days, and then placed under light for 7 days. **B)** Survival rates of seedlings in panel A. Seedlings remaining green were considered alive. Data represent mean ± SEM from three biological replicates. At least 45 seedlings were included per replicate, * for *p*<0.05 (Student’s t-test).

### UPL3 is a Negative Regulator of BR Response with a Role in Growth Attenuation under Starvation Conditions

We obtained two T-DNA insertion alleles of *upl3* mutant (*upl3-4* and *upl3-26*) and analyzed their phenotypes. The *upl3-4* mutant, in which the T-DNA is inserted in the 8^th^ exon, is reported to be a UPL3 null-allele (Furniss et al., 2018). The *upl3-26* allele contains a T-DNA inserted in the 15^th^ intron and expresses a decreased level of *UPL3* RNA (Figure 2A and S4A), and thus is expected to be a weak allele. Indeed, *upl3-4* displayed stronger phenotypes than *upl3-26*. When grown in soil under light, the *upl3* mutants had elongated petiole and narrow leaf blades, similar to the phenotype of transgenic plants overexpressing BRI1-GFP, compared to the wild-type plants (Figure 2B). In contrast, the *baf1* mutant showed no obvious difference in morphology from the wild type (Figures 2B and S4B). We quantified the phenotypes by measuring the length/width ratio of leaves (Figure 2C). The average length/width ratios of *upl3-4, BRI1-GFP,* and *baf1* mutants were 58%, 78%, and 8%, respectively, bigger than that of wild type (Figures 2C). In addition, the hypocotyl elongation of *upl3* mutants showed a lower sensitivity to propiconazole (PPZ, a BR biosynthesis inhibitor) than wild type and *baf1*, whereas *baf1* is slightly less sensitive to PPZ than the wild type (Figures 2D and 2E). These phenotypes indicate that UPL3 and BAF1 are negative regulators of BR response, with UPL3 playing a more prominent role than BAF1.

To test whether UPL3 plays a role in sugar regulation of BR signaling, we grew the seedlings on sugar-containing or sugar-free media under light for 4 days and then shifted them into darkness for 3 days. The growth of wild-type seedlings on sugar-free medium is attenuated due to depletion of photosynthate and starvation-induced degradation of BZR1 (Zhang et al., 2016). The *upl3* mutants displayed similar morphology as the wild-type on sugar-containing media but longer hypocotyls than wild-type on sugar-free media (Figure 2F and 2G), suggesting that UPL3 is required for growth arrest under starvation conditions. In contrast to the *upl3* mutants, the *baf1* mutant showed a shorter hypocotyl than wild-type under sugar starvation conditions (Figure 2F and 2G).

### UPL3 Plays a Prominent Role in Supporting Plant Survival from Starvation

In order to test the functions of UPL3 and BAF1 in plant survival of sugar starvation, we perform long-term sugar starvation assays. We grew *upl3* and *baf1* mutants together with some BR mutants on sugar-free media under light for 4 days, transferred them into the dark for 9 to 12 days, and then let them recover under light for 7 days. The numbers of dead and alive seedlings were counted (Figure 3). Increasing dark periods caused seedling death in all genotypes, presumably due to sugar starvation resulting from the absence of photosynthesis. Compared to the wild type, the survival rate was increased in the BR deficient mutant *det2* but decreased in the *bzr1-1D* mutant, transgenic plants overexpressing BRI1, and *upl3* mutants. The *upl3-4* mutant showed a similarly severe phenotype as *bzr1-1D*, whereas the phenotype of *upl3-26* was weaker. The survival rate of *baf1* was slightly lower than the wild type but much higher than *upl3-4* (Figures 3A and 3B). These results show that UPL3 plays a prominent role in plant survival of sugar starvation.

### UPL3 is Required for the Starvation-induced Degradation of BZR1

We analyzed the BZR1 protein level using an anti-BZR1 antibody. The *upl3-26* mutant accumulated much higher levels of both phosphorylated and dephosphorylated forms of BZR1 than the wild type (Figure 4A). We also crossed a *BZR1:BZR1-CFP/*Col-0 transgenic line with *upl3-26*, and found more BZR1-CFP protein accumulated in *upl3-26* than in the wild-type background (Figure 4B). The RNA levels of BZR1-CFP are similar in different backgrounds (Figure 4C), suggesting that UPL3 promotes the degradation of BZR1. To test whether UPL3 is involved in the starvation-induced degradation of BZR1, we analyzed the BZR1 protein levels by immunoblotting in plants grown under the sugar-free and sugar-supplemented conditions as in Figures 2F and 2G. We observed higher levels of both phosphorylated and dephosphorylated BZR1 in *upl3-26* than in wild type grown on sugar-free media, but similar BZR1 protein levels between *upl3-26* and wild type grown on sugar-containing media (Figure 4D). The results indicate that UPL3 plays a specific role in starvation-induced BZR1 degradation.

**Figure 4.**
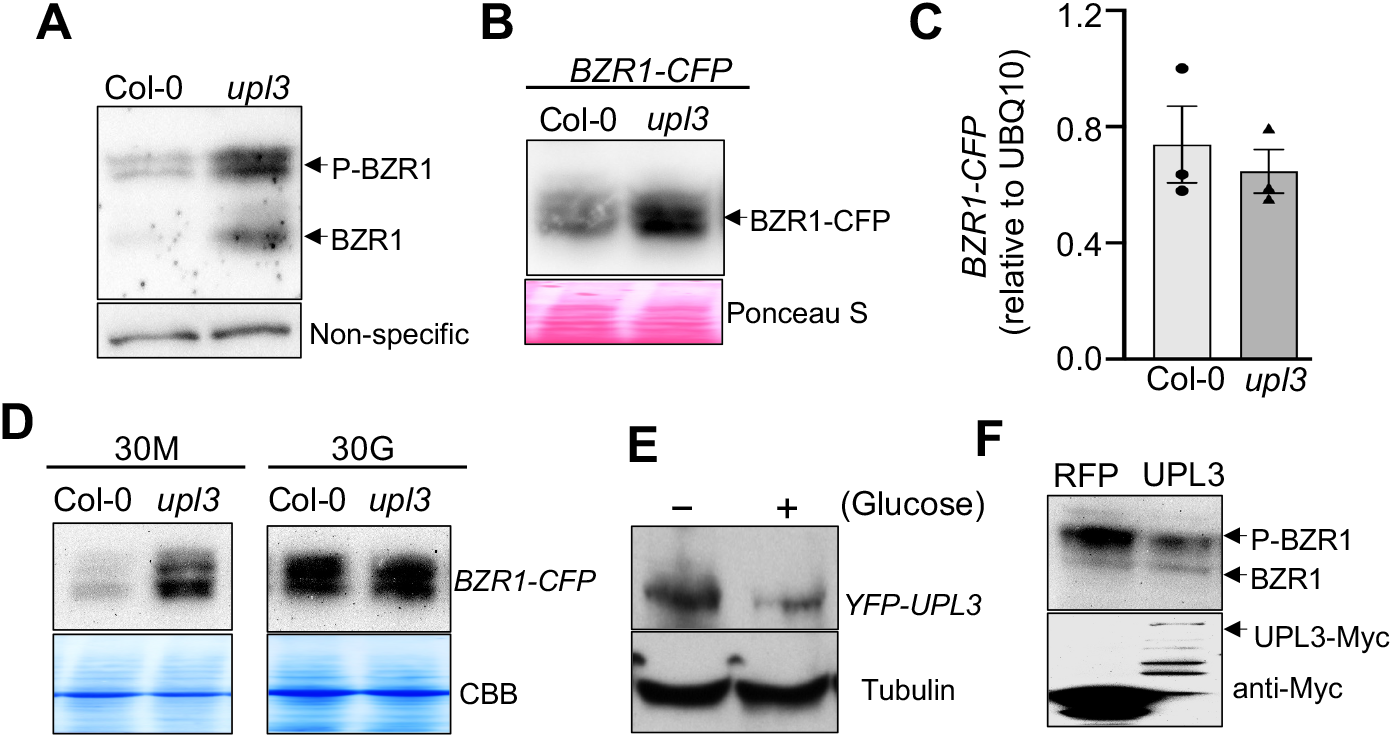
UPL3 mediates sugar regulation of BZR1 degradation. **A)** Immunoblot shows BZR1 protein accumulation in *upl3-26* mutants grown in the dark for 5 days. **B-D)** BZR1 protein accumulates in *upl3* mutant under sugar starvation conditions. The *BZR1:BZR1-CFP* transgene was crossed from Col-0 into *upl3-26* background. Seedlings were grown under constant light and then treated by 48-hr darkness (B and C) or grown under light and then in the dark for 3 days on media containing 30 mM mannitol (30M) or 30 mM glucose (30G) (D). The samples were analyzed by anti-GFP immunoblot (B and D) or qRT-PCR (C). Ponceau S and Coomassie Brilliant Blue (CBB) staining show the loading. **E)** Sugar decreases the UPL3 protein level. The *35S:YFP-UPL3/upl3-4* seedlings were grown in sugar-free liquid media and treated with 15 mM glucose for 2 hours, and then analyzed by immunoblotting using anti-GFP or anti-tubulin antibodies. **F)** UPL3 promotes BZR1 degradation. RFP-Myc or UPL3-Myc was transiently co-expressed with BZR1-YFP in *N. benthamiana.* The proteins were analyzed by immunoblotting with anti-GFP and anti-Myc antibodies.

We then tested whether starvation alters the UPL3 protein level. We grew the *35S:YFP-UPL3/upl3-4* transgenic plants in sugar-free liquid media for eight days and then added glucose to the culture media for 2 hours. The glucose treatment significantly decreased the YFP-UPL3 level (Figure 4E), indicating that sugar reduces UPL3 level through a posttranscriptional mechanism. To test the effect of UPL3 overexpression on BZR1 accumulation, we co-expressed BZR1-YFP with UPL3-Myc in *Nicotiana benthamiana* leaves. As shown in Figure 4F, UPL3-myc reduced the BZR1-YFP protein level, indicating that an elevated level of UPL3 promotes the degradation of BZR1.

### UPL3 Mediates Sugar-TOR Regulation and the Autophagy-dependent BZR1 Degradation

Previous studies showed that the starvation-induced BZR1 degradation is regulated by the TOR signaling pathway and mediated by both 26S-proteasome pathway or autophagy pathway (Kim et al., 2014; Zhang et al., 2016; Nolan et al., 2017b; Kim et al., 2019; Wang et al., 2021). We therefore analyzed the functions of UPL3 in these specific processes. The inhibitor of TOR, AZD8055, caused BZR1 degradation in the wild-type, but the effect is much weaker in *upl3-26* (Figure 5A), suggesting that UPL3 contributes to the BZR1 degradation caused by inactivation of TOR. We then tested the effects of inhibitors of proteasome (MG132) and autophagy (3-MA) on BZR1 accumulation in wild-type and *upl3-26* background. In three independent experiments, the abundance of BZR1 protein was about 3.3x that of the wild-type without inhibitor treatment. MG132 treatment increased BZR1 protein level by about 2.4x and 1.7x in wild type and *upl3* mutant background, respectively. In contrast, 3-MA treatment increased BZR1 level by 1.6x in wild-type background but had no obvious effects on BZR1 level in *upl3* (Figures 5B-5C). The results indicate that UPL3 is required for autophagy-dependent BZR1 degradation.

**Figure 5.**
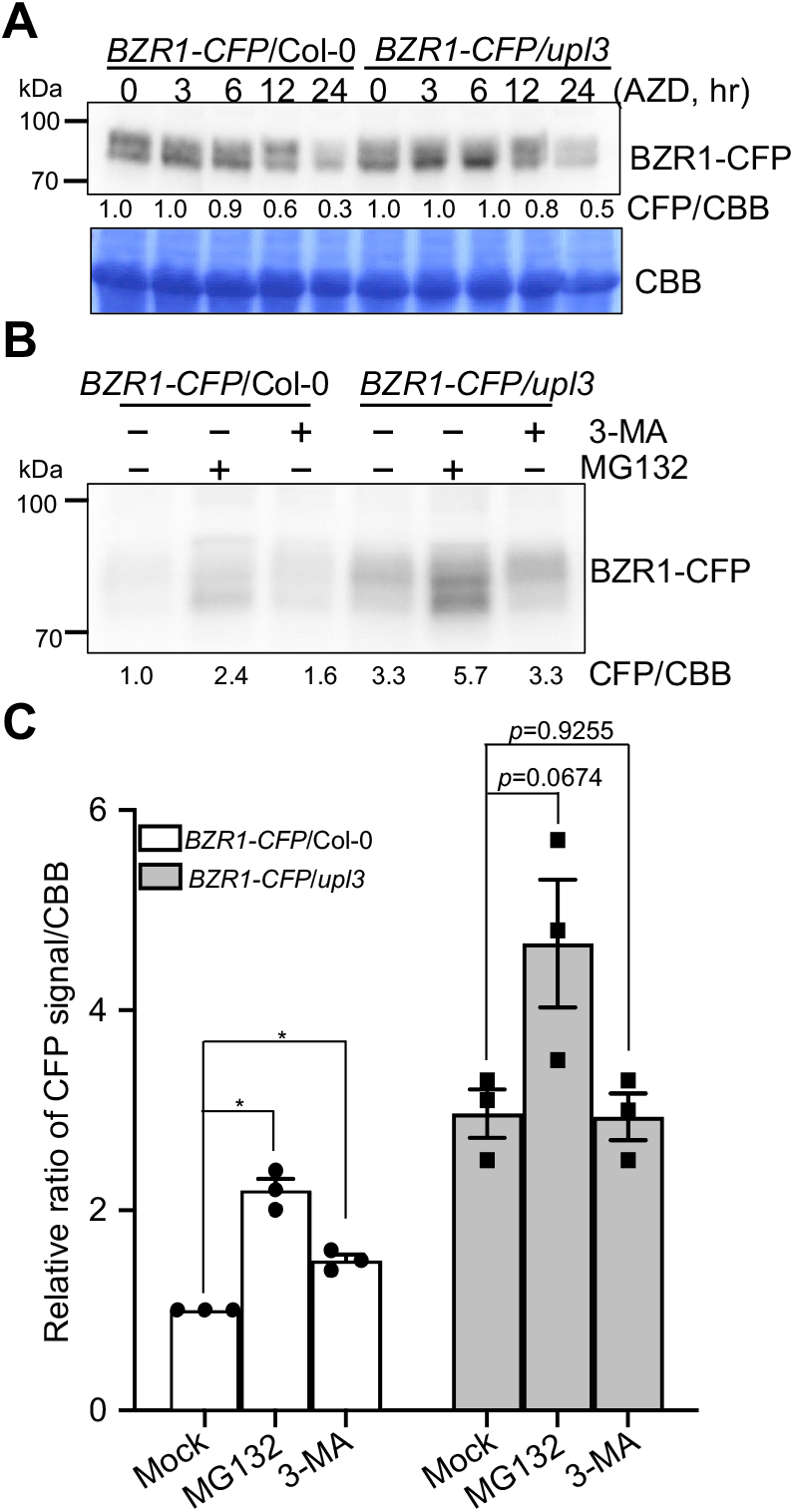
UPL3 is required for autophagy-dependent BZR1 degradation. **A)** UPL3 is involved in BZR1 degradation caused by TOR inactivation. The *BZR1-CFP/Col-0* and *BZR1-CFP/upl3-26* seedlings were grown in liquid media containing 30 mM glucose for 5 days and then treated with 5 μM AZD-8055 or DMSO (mock) for the indicated times. BZR1-CFP was analyzed by anti-GFP immunoblotting. Numbers below blots are band intensities relative to the mock treatments. Coomassie Brilliant Blue (CBB) staining of the blot shows the loading. **B)** UPL3 is required for autophagy-dependent BZR1 degradation. The *BZR1-CFP/Col-0* and *BZR1-CFP/upl3-26* seedlings grew on 1/2 MS medium containing 30 mM mannitol under light for 4 days and in the dark for 12 hrs. MG-132 (30 μM), 3-MA (5 mM), or mock solution was added to the media. The plants were kept in the dark for 36 hours and then analyzed by anti-GFP immunoblotting. The numbers below the gel image are relative band intensity. **C)** The relative signal intensity normalized to the intensity of CBB staining. Data are mean ± SEM., *n* = 3 independent experiments. * for *p*<0.05.

## DISCUSSION

The crosstalk between sugar-sensing and growth hormone pathways is crucial for optimal growth and survival under changing environmental conditions. BR signaling is essential for maximum growth using available photosynthates. However, when the levels of photosynthates are low, such as when plants are shaded or stressed, the BR signaling output must be attenuated to avoid cellular starvation and to improve survival. Therefore, understanding the molecular mechanisms of sugar-BR crosstalk is important for improving plant productivity and resilience.

Previous studies by us and other researchers have shown that sugar signaling through the TOR pathway regulates the degradation of BZR1 and its homolog BZR2/BES1 through autophagy-dependent pathways. The degradation of BZR1 and BZR2/BES1 under starvation conditions contributes to growth arrest, which alleviates cellular starvation (Zhang et al., 2016; Nolan et al., 2017b). Here, we identified the BZR1-interacting proteins that mediate the starvation-induced BZR1 degradation. Our genetic and biochemical evidence indicates that UPL3 interacts with BZR1 to promote BZR1 degradation, causing growth arrest and increasing plant survival under sugar starvation conditions. Our study also shows that BZR1 interacts with the F-box E3 ubiquitin ligase BAF1, which was reported recently as an E3 ligase for BZR2/BES1 (Wang et al., 2021). The stronger phenotypes of *upl3* than *baf1* indicate that UPL3 plays a more important role than BAF1.

Identification of *in vivo* interacting proteins that mediate the degradation of a target protein of interest is likely to be challenging because the interaction leads to the degradation and reduced level of the bait protein. Further, the protein that mediates starvation-induced BZR1 degradation is likely to interact with BZR1 only under sugar starvation conditions. Therefore, we chose to use BZR1-overexpressing transgenic plants grown under starvation conditions to perform IP-MS and we also used metabolic stable isotope labeling followed by immunoprecipitation and mass spectrometry (SILIP-MS) to quantitatively distinguish between BZR1-binding proteins and non-specific background. The experiment identified 28 known BZR1 interactors and their family members and 41 putative new BZR1 interactors that showed similar enrichment as the known interactors. While some of the newly identified interactors might represent artifacts due to BZR1 overexpression, our follow-up genetic characterization confirmed the function of UPL3 in BZR1 degradation, growth arrest, and survival under starvation conditions.

Specifically, a series of experiments confirmed UPL3’s function in starvation-induced BZR1 degradation. First, the UPL3-BZR1 interaction was confirmed by immunoprecipitation using targeted mass spectrometry and by co-IP (Figure 1). Second, the loss-of-function *upl3* mutants accumulate increased levels of BZR1 protein when grown without the supplement of sugar but accumulate a normal level of BZR1 when grown on a medium containing sugar (Figure 4D), indicating a specific function of UPL3 in starvation-induced degradation of BZR1. Third, starvation increased UPL3 protein level and overexpression of UPL3 reduced the accumulation of BZR1 (Figure 4E-F).

Consistent with the accumulation of BZR1, the *upl3* mutants display a range of phenotypes that resemble plants overexpressing BRI1. These phenotypes include long petioles and slim leaves when grown under light, long-hypocotyls when grown on BR inhibitor in the dark or grown on sugar-free medium and transferred from light to dark (Figure 2), and reduced survival after a long time in the dark (starvation) (Figure 3). Importantly, the *upl3* mutants showed increases in BZR1 protein level and hypocotyl length under starvation conditions, but accumulated a normal level of BZR1 protein and showed normal hypocotyl length compared to the wild type when grown on a sugar-containing medium. Together, these results demonstrate that UPL3 plays a role in BZR1 degradation specifically under starvation condition.

UPL3 is a member of the HECT (Homologous to E6-AP Carboxyl Terminus) family ubiquitin E3 ligases, which are proteasome-associated E3 ligases. HECT E3s accept ubiquitin from an E2 and covalently link it to a conserved Cys in their HECT domain before transferring ubiquitin to the protein substrate (Rotin and Kumar, 2009). The Arabidopsis genome encodes seven HECT E3 ligases that are grouped into four subfamilies (UPL1/2, UPL3/4, UPL5, and UPL6/7) (Downes et al., 2003). The UPL3 and UPL4 subfamily encodes approximately 200-kDs proteins containing C-terminal HECT domain and N-terminal Armadillo repeads. The *upl3* (*kaktus*) mutants have been reported to show a wide range of phenotypes, including hyperbranched trichome with increased endoreduplication (Downes et al., 2003; El Refy et al., 2003), hypersensitivity to gibberellin (GA) (Downes et al., 2003), increased seed size (Miller et al., 2019), defects in salicylic acid (SA)-induced gene expression and pathogen defense (Furniss et al., 2018). The *upl3upl4* double mutants have severe growth defect and enhanced ethylene response (Wang et al., 2022). Several target proteins of UPL3 have been identified to explain some of these phenotypes. These include GLABROUS 3 (GL3) and ENHANCER OF GLABROUS 3 (EGL3) transcription factors and the cyclin-dependent kinse inhibitor KRP2 in trichome development (Patra et al., 2013; Xue et al., 2023); the LEC2 transcription factor in regulating seed development (Miller et al., 2019), NPR1 in SA signaling and immunity (Wang et al., 2022), and the ethylene response factor ETHYLENE INSENSITIVE 3 (EIN3) (Wang et al., 2022). Our observation of BZR1 accumulation in *upl3* may explain the GA-hypersensitivity phenotype, as BZR1 is an essential target of the GA-signaling DELLA repressors (Bai et al., 2012).

Recent studies suggested that UPL3 acts downstream of pathway-specific E3 ubiquitin ligases to together regulate the activity and degradation of hormone-responsive transcription factors NPR1 and EIN3 (Wang et al., 2022). The defect of *upl3* mutant in SA-induced gene expression and immunity is associated with complex effects on the translation, polyubiquitination, and degradation of the SA-response factor NPR1. The ubiquitination of NPR1 appears to be a progressive event in which initial modification by a Cullin-RING E3 ligase promotes its chromatin association and expression of target genes. Subsequent polyubiquitination of NPR1 by the E4 ligase, UBE4, inhibits NPR1 transcriptional activity. The relay of NPR1 from UBE4 to UPL3 and its homologs leads to further polyubiquitination and degradation of inactive NPR1. Similarly, there is evidence that UPL3 and its homolog UPL4 mediate polyubiquitination and degradation of EIN3 following the initial ubiquitination of EIN3 by the F-box E3 ligase EBF1/2. Further, the *in vivo* interaction between UPL3 and EIN3 requires EBF2 (Wang et al., 2022). The studies suggest that these SA- and ethylene-responsive TFs are relayed from pathway-specific ubiquitin ligases to proteasome-associated UPLs, and further such ubiquitin ligase relay may be a universal mechanism for UPL functions (Wang et al., 2022). It is possible that UPL3 acts in relay with other E3 ligases in BZR1 degradation.

BZR1 and its close homolog BZR2/BES1 are targets of several E3 ligases, including PUB40 (Kim et al., 2019), SINAT (Yang et al., 2017), and BAF1 (Wang et al., 2021). Our BZR1 IP-MS analysis under the dark-induced starvation condition detected BAF1 but not SINAT or PUB40, supporting the role of BAF1 in starvation-induced BZR1 degradation. BAF1 was reported recently as an E3 ligase that interacts with BZR2/BES1 (Wang et al., 2021). Interestingly, the phenotypes of *baf1* mutant is distinct from those of *upl3*. When grown under normal conditions in soil or on solid media under light, *baf1* showed no obvious phenotype (Figure 2) (Wang et al., 2021). The *baf1* seedlings were slightly less sensitive to PPZ than the wild type but more sensitive than the *upl3* mutants (Figure 2E). Surprisingly, when light-grown seedlings were transferred into darkness without exogenous sugar, *baf1* showed shorter hypocotyls than wild type (Figure 2F-G), raising the possibility that BAF1 positively regulates BZR1/2 activity under starvation conditions. After long-term starvation in the dark, the survival rate of *baf1* mutant was slightly lower than the wild type but much higher than *upl3* (Figure 3). These observations indicate that UPL3 plays a more prominent role than BAF1 in the starvation-induced degradation of BZR1. It is possible that, like NPR1 and EIN3 (Wang et al., 2022), the transcriptional activity and degradation of BZR1/BES1 are regulated by progressive unbiquitination: monoubiquitination or limited polyubiquitination by BAF1 increases BZR1/BES1 transcriptional activity while leading to polyubiquitination by UPL3, which promotes BZR1/BES1 degradation by the autophagy pathway. Although UPLs are considered proteasome-associated E3 ligases, a large body of evidence supports the emerging role of HECT E3 ligases in autophagy (Melino et al., 2019).

Our study reveals the role of UPL3 in sugar regulation of plant growth and survival by mediating starvation-induced BZR1 degradation and gating BR signaling outputs. We also observed sugar affecting the level of UPL3 protein in a posttranscriptional manner (Figure 4E), suggesting a central role for UPL3 in mediating broader sugar responses. Therefore, whether and how UPL3 mediates sugar regulation of its additional target proteins and their diverse downstream responses will be interesting questions to be investigated in the future. It’s intriguing to notice that proteomic study revealed that UPL3 plays a role in the ubiquitination and degradation of the sugar sensor HEXOKINASE 1 (HXK1) and many enzymes involved in carbohydrate metabolism (Lan et al., 2022). Together, the evidence suggests that UPL3 may play a central role in metabolic regulation of growth and environmental adaptation.

## EXPERIMENTAL PROCEDURES

### Plant Materials and Growth Conditions

The *upl3-26* T-DNA insertional mutant (SALK_116326C,) was obtained from the Arabidopsis Biological Resource Center. The *baf1* mutant (GABI_001F08) was the same allele as reported (Wang et al., 2021) and obtained from GABI-Kat database. The *upl3-4*, *35S:YFP-UPL3*/*upl3* (Furniss et al., 2018) seeds were kindly provided by Dr. Steven H. Spoel. Genotyping was performed with primers listed in Supplemental Table S1 using the Phire Tissue Direct PCR Master Mix (Thermo Fisher, #F170L). The BR-related mutants *bzr1-1D*, *det2* (Wang et al., 2002), *BZR1-CFP*/Col-0 (He et al., 2005), *BZR1-YFP*/Col-0 (Gendron et al., 2012), and *BRI1-GFP* (Wang et al., 2001) were reported previously. All the plants were in Col-0 ecotype background. Plants were all grown in a greenhouse with a 16-hr light/8-hr dark cycle at 22-24℃ for general growth, phenotype observation, and seed harvesting.

### Plasmid Constructs

The full-length coding sequences of UPL3 or BZR1 without the stop codons were cloned into pENTR/SD/D-TOPO plasmid. Then the UPL3 TOPO or BZR1 TOPO entry was inserted into the modified 35S-gcpCAMBIA1390-4myc-6His vector or pEarleyGate 101 vector to generate the 35S:UPL3-myc or 35S:BZR1-YFP construct. All the primers used in this study are listed in Supplemental Table S1.

### Seedling phenotype analysis

For hypocotyl phynotype analysis, seeds were sterilized by 75% (v/v) ethanol and sown on solid agar plates (0.8% phytoblend) containing half-strength Murashige and Skoog medium (1/2 MS) (pH 5.7) and sugar or hormone inhibitors as designed. After three days of incubation at 4℃ and seeds were grown in a growth chember at 22℃ under donstent light or darkness. For starvation treatment, plants were grown under light for 4 days and then in the dark for various times. Pictures were taken and the hypocotyl lengths were measured using the Image J software.

For starvation survival assays, seedlings were grown on sugar-free medium under constant light for 4 days, put in the dark for 9 to 12 days, and back to light for 7 days. Pictures were taken and survival rates were calculated. Seedlings remaining green were considered alive. At least 45 seedlings were included in each of the three replicates.

### Tobacco Infiltration and Coimmunoprecipitation

Agrobacterium (GV3101) cells harboring the construct expressing BZR1-YFP, 35S:YFP, or UPL3-Myc were resuspended in the infiltration medium (10 mM of MES at pH 5.6, 10 mM of MgCl2, 0.01% tween 20 and 150 mM of acetosyringone), mixed as indicated, and infiltrated into leaves of *Nicotiana benthamiana*. Tissues were harvest 36 hrs later for coimmunoprecipitation (Co-IP) and immunoblot analysis.

After grinding with liquid nitrogen, the tissues were resuspended in immunoprecipitation (IP) extraction buffer (50 mM Tris-HCl, pH 7.5, 50 mM NaCl, 10% glycerol, 0.25% Triton X-100, 0.25% NP40,1 mM PMSF, Pierce protease inhibitor cocktail [Thermo Fisher Scientific], and PhosStop cocktail [Roche]), and sonicated for 20 seconds (1s on/off). After centrifugation at 14,000 rpm for 15 min, the supernatant was incubated with Anti-GFP Magarose Beads (Smart-Lifesciences, SM038005) for 1 hr in the cold room. The beads were washed three times with 1 mL of IP buffer without detergent. The proteins were eluted with 50 μl SDS sample buffer and separated by SDS-PAGE.

### Chemical Inhibitor Treatments

Seedlings were grown in liquid 1/2 MS media with 30 mM glucose for 5 days and then treated with 5 μM AZD-8055 (Selleck, S1555) or DMSO (mock) for indicated hours. Tissue was harvested for immunoblotting.

Seedlings were grown on 1/2 MS medium with 30 mM mannitol under light for 4 days and placed in the dark. After 12 h in the dark, MG-132 (Millipore, 203790) or 3-MA (Selleck, S2767) were added to the media (final concentration 30 μM and 5 mM, respectively) under green safe light, and tissues were kept in the dark for 36 h before harvested for immunoblotting.

### Protein Extraction and Immunoblot Analysis

For protein extraction, plants were frozen in liquid nitrogen, ground, weighed, and added into corresponding 2× SDS buffer (0.125 mM Tris-HCl [pH 6.8], 4% SDS, 20% Glycerol and 2% β-mercapto-ethanol). Samples were heated for 10 min at 95°C, centrifuged at 10000g for 10 min, separated on a 7.5% acrylamide gel and then blotted on PVDF membranes (Millipore, IPVH0010) in 192 mM glycine and 25 mM Tris-HCl. Membranes were blocked for 1 hour at room temperature in a blotting buffer (140 mM NaCl, 10 mM KCl, 8 mM Na_2_HPO4, 2 mM KH_2_PO4, 0.5% skim milk, and 0.1% Tween20, pH 7.4). The gel blots were incubated with the primary antibodies (anti-GFP, Transgen Biotech, HT801, at 1:1000 dilution; anti-BZR1, custom-made, at 1 µg/ml; anti-Actin, Sigma, A2228, at 1:5000; anti-Tubulin, Abmart, M20023, at 1:5000 dilutions) for 1 hr at room temperature or overnight in the cold room. The secondary antibodies were used at 1:5000 dilutions for 1 hour.

### ^15^N Stable Isotope Labeling for Quantitative MS Analysis of the BZR1 interactome

The *35S:BZR1-YFP* and the *35S:GFP* plants were grown for 2 weeks at 22°C under constant light on vertical plates containing ^14^N or ^15^N medium (half-strength, 0.39 g/L) Murashige and Skoog modified basal salt mixture without nitrogen, 8 g/L phytoblend, and 0.5 g/L KNO3, 0.5 g/L NH_4_NO3 or 0.5 g/L K^15^NO3, 0.5 g/L ^15^NH_4_^15^NO3 [Cambridge Isotope Laboratories], pH 5.8). The plants were put in darkness for 24 hrs and then harvested and ground in liquid nitrogen. Proteins were extracted from 1 g of tissue in 2 mL of IP buffer (50 mM Tris-HCl, pH 7.5, 50 mM NaCl, 10% glycerol, 0.25% Triton X-100, 0.25% NP40, 1 mM PMSF, Pierce protease inhibitor cocktail [Thermo Fisher Scientific], and PhosStop cocktail [Roche]). After centrifugation at 14,000 rpm for 15 min, the supernatant was incubated with GFP-trap (Anti-GFP Magarose Beads, Smart-Lifesciences, SM038005) for 1 hr in the cold room. The beads were washed three times with 1 mL of IP buffer without detergent. The beads of ^14^N-labeled BZR1-YFP and the ^15^N-labeled 35S:GFP samples (or ^14^N-labeled 35S:GFP and ^15^N-labeled 35S:BZR1-YFP) were combined. The proteins were eluted with 50 μl SDS buffer and separated by SDS-PAGE.

Following Coomassie Brilliant Blue staining, the protein bands were cut and were washed three with 50% acetonitrile in 25 mm ammonium bicarbonate (NH_4_HCO_3_) and vacuum-dried. The gel samples were next reduced with DTT (10 mm in 25 mm NH_4_HCO_3_ at 56 °C for 1 h) and alkylated with iodoacetamide (50 mm in 25 mm NH_4_HCO_3_ at room temperature for 45 min). Then the gel pieces were vacuum-dried, rehydrated in 5-10 μl of digestion buffer based on the gel size (10 ng/μl trypsin in 25 mm NH_4_HCO_3_), and covered with a minimum volume of NH_4_HCO_3_. After overnight digestion at 37 °C, peptides were extracted twice with a solution containing 50% acetonitrile and 0.1% formic acid. The extracted digests were vacuum-dried and resuspended in 5 μl of 0.1% formic acid in water. Peptides are desalted by C18 pipette tip.

Peptides were analyzed by liquid chromatography–tandem mass spectrometry (LC-MS) on an Easy LC 1200 UPLC liquid chromatography system (Thermo Fisher) connected to Orbitrap Q Exactive HF (Thermo Fisher) for DDA analysis or connected to Orbitrap Eclipse (Thermo Fisher) for targeted quantification (PRM). Peptides were first trapped using trapping column Acclaim PepMap 100 (75 µm x 2cm, nanoViper 2Pk, C18, 3 µm, 100A), then separated using analytical column AUR2-25075C18A, 25CM Aurora Series Emitter Column (25cm x75 µm, 1.6um C18) (IonOpticks). The flow rate was 300 nL/min, and a 120 min gradient was used. Peptides were eluted by a gradient from 3 to 28% solvent B (80% (v/v) acetonitrile/0.1% (v/v) formic acid) over 100 min and from 28 to 44% solvent B over 20 min, followed by a short wash at 90% solvent B. Precursor scan was from mass-to-charge ratio (m/z) 375 to 1600 (resolution 120,000; AGC 3.0E6) and top 20 most intense multiply charged precursors were selected for fragmentation. Peptides were fragmented with higher-energy collision dissociation (HCD) with normalized collision energy (NCE) 27. For targeted quantification, data was acquired in the PRM mode. Peptides were separated using an EasyLC1200 system (Thermo) connected to a high-performance quadrupole Orbitrap mass spectrometer Eclipse (Thermo). The LC setting was the same as described in DDA. The samples were analyzed using PRM mode with an isolation window of 1.6 Th. PRM scans were done using 30,000 resolutions (AGC target 1.5 e5, 300% AGC) triggered by an inclusion list. Normalized collision energy of 27% was used in a higher-energy dissociation mode.

MS/MS data were converted to peaklist using a script PAVA (peaklist generator that provides centroid MS2 peaklist (Guan et al., 2011; Shrestha et al., 2022), and data were searched using Protein Prospector against the TAIR database Arabidopsis thaliana from December 2010 (https://www.arabidopsis.org/), concatenated with sequence randomized versions of each protein (a total of 35386 entries). A precursor mass tolerance was set to 10 ppm and MS/MS2 tolerance was set to 20 ppm. Carbamidomethyl was searched as a constant modification. Variable modifications include protein N-terminal acetylation, peptide N-terminal Gln conversion to pyroglutamate, Met oxidation. FDR 1% was set for both proteins and peptides. The cleavage specificity was set to trypsin, allowing two missed cleavage and a maximum of two modifications. For quantification, ^15^N labeling efficiency was manually checked. ^15^N labeling’ was chosen as a quantitative method using Protein Prospector with automatic adjustment of L:H intensity ratios with labeling efficiency. Quantification of DDA data was done as described (Shrestha et al., 2022). Quantification of PRM data using Skyline was done as described previously (Reyes et al., 2022).

### Accession Numbers

BZR1(AT1G75080), UPL3 (AT4G38600), BAF1(AT1G76920)

### Supplemental Data

Supplemental Figure S1. Identify BZR1-interacting proteins by quantitative proteomics.

Supplemental Figure S2. Quantitation of enrichment in BZR1-YFP relative to 35S:GFP control in SILIP-MS.

Supplemental Figure S3. Targeted quantification of UPL3 and ACC1 in three BZR1 SILIP-MS experiments..

Supplemental Figure S4. Characterization of the *upl3-26* and *baf1* mutants. Supplemental Table S1. Primers used in this study

Supplemental Data Set S1. SILIP-MS analysis of BZR1 interactors.

### Data availability

The mass spectrometry proteomics data have been deposited to the ProteomeXchange Consortium via the PRIDE (Perez-Riverol et al., 2022) partner repository with the dataset identifier PXD04613.

The author(s) responsible for distribution of materials integral to the findings presented in this article in accordance with the policy described in the Instructions for Authors (https://academic.oup.com/plcell/pages/General-Instructions) is : Zhi-yong Wang

## Acknowledgments

We thank Dr. Steven H. Spoel for the *upl3-4* and 35S:*YFP-UPL3/upl3-4* seeds. We also thank Ruben Shrestha and Andres V Reyes for helping in MS data acquisition and analysis.

## Funding and additional information

The work was supported by grants from the National Institute of Health (NIH, R01GM066258 to Z-Y.W., NIH, S10OD030441 to S.-L.X.), the National Natural Science Foundation of China grant (NO. 31800239, http://www.nsfc.gov.cn) and the China Scholarship Council to Z.Z. The funders had no role in study design, data collection and analysis, decision to publish, or preparation of the manuscript. The work was also funded by the Carnegie endowment fund to the Carnegie Mass Spectrometry Facility.

## Author contribution

Z.Z., and Z.-Y.W. designed experiments, analyzed data, and wrote the manuscript. Z.Z performed all experiments except those specified below. H.Z. performed the experiments on starvation survival, verification of UPL3-BZR1 interaction, and effects of TOR and autophagy inhibitors on BZR1. E.G. made the constructions of UPL3-Myc and did mutant genotyping. T.G and S.X performed PRM-MS and analyzed the data.

---

**Figure S1.**
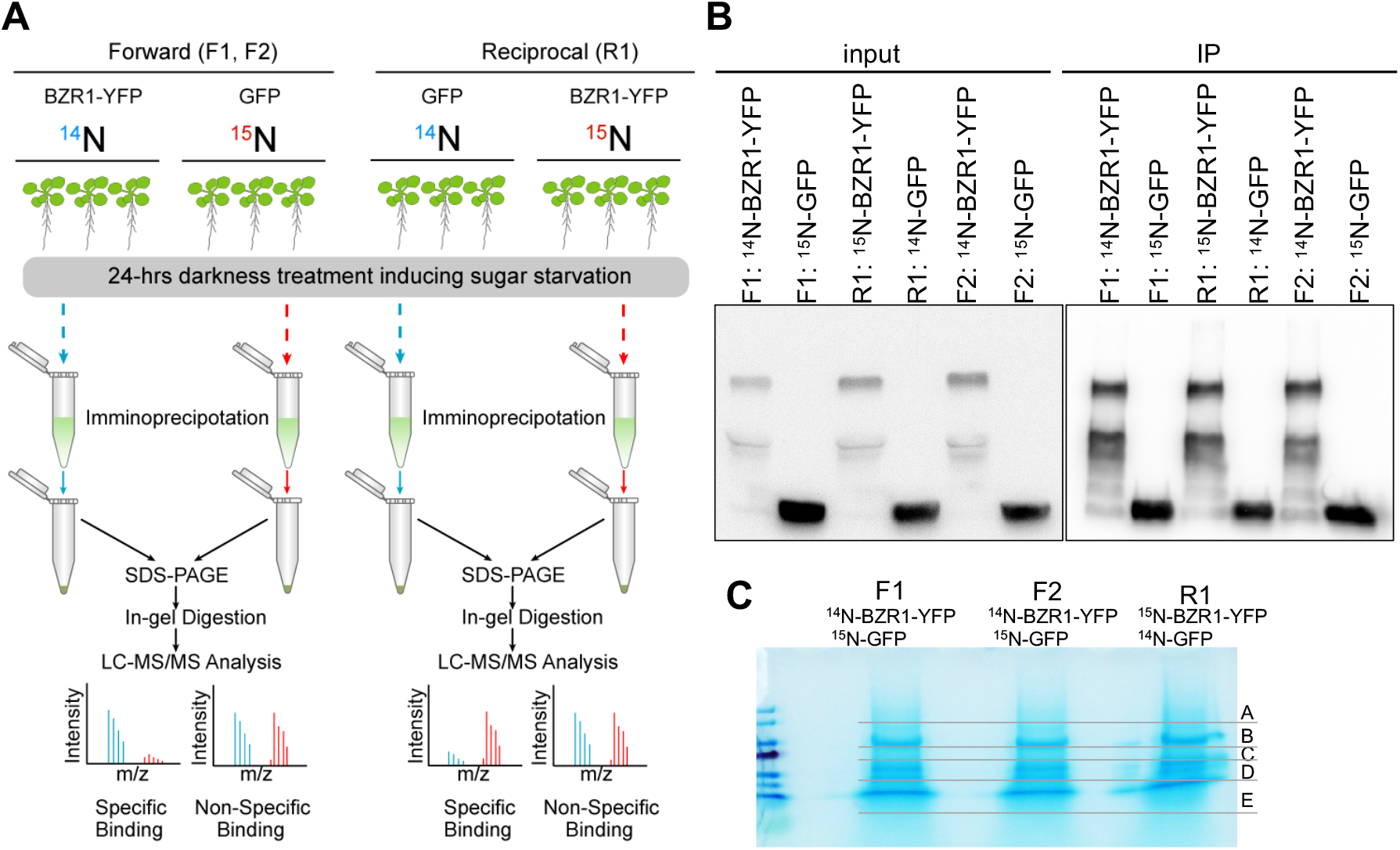
Identify BZR1-interacting proteins by quantitative proteomics. **A)** Workflow of SILIP-MS experiments for identifying BZR1-interacting proteins in sugar-starved Arabidopsis plants. Two-week seedlings grown on ^14^N- or ^15^N-labeled 1/2MS medium without sugar were treated by darkness for 24 hrs, then were harvested for immunoprecipitation. In two forward-labeling replicate experiments (F1 and F2), the *35S:BZR1-YFP* and *35S:GFP* plants were labeled with ^14^N and ^15^N, respectively, whereas the isotopes were switched in the reverse labeling replicate (R1). **B)** Western blots show the *BZR1-YFP* or *GFP* protein immunoprecipitated by GFP-trap. **C)** Coomassie blue-stained SDS-PAGE gels of proteins immunoprecipitated by GFP-trap from ^14^N/^15^N-labelled *BZR1-YFP* or *GFP*. Each lane was cut into 5 pieces as shown in the image for mass spectrometry analysis.

**Figure S2.**
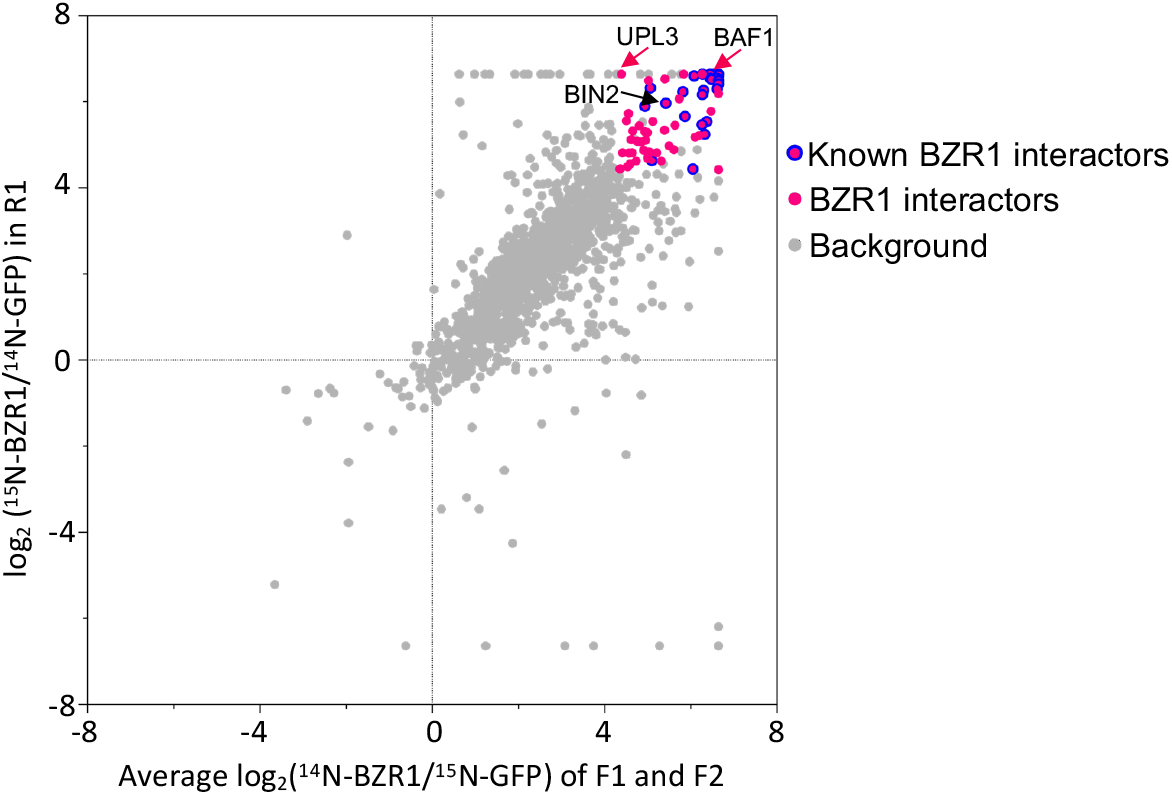
Quantitation of enrichment in BZR1-YFP relative to 35S:GFP control in SILIP-MS. A plot of the signal ratios between *BZR1-YFP* and *GFP* for proteins detected in three replicate experiments (average of F1 and F2 *vs* R1). The pink dots indicate proteins with log2(BZR1/GFP) >=4.34 and at least two peptides identified in each replicate. The pink dots with blue border indicate the reported BZR1 interactors and their family members. The light gray dots indicate proteins detected by SILIP-MS with fold change log2(BZR1/GFP) <4.34 or less than two peptides identified in any replicate.

**Figure S3.**
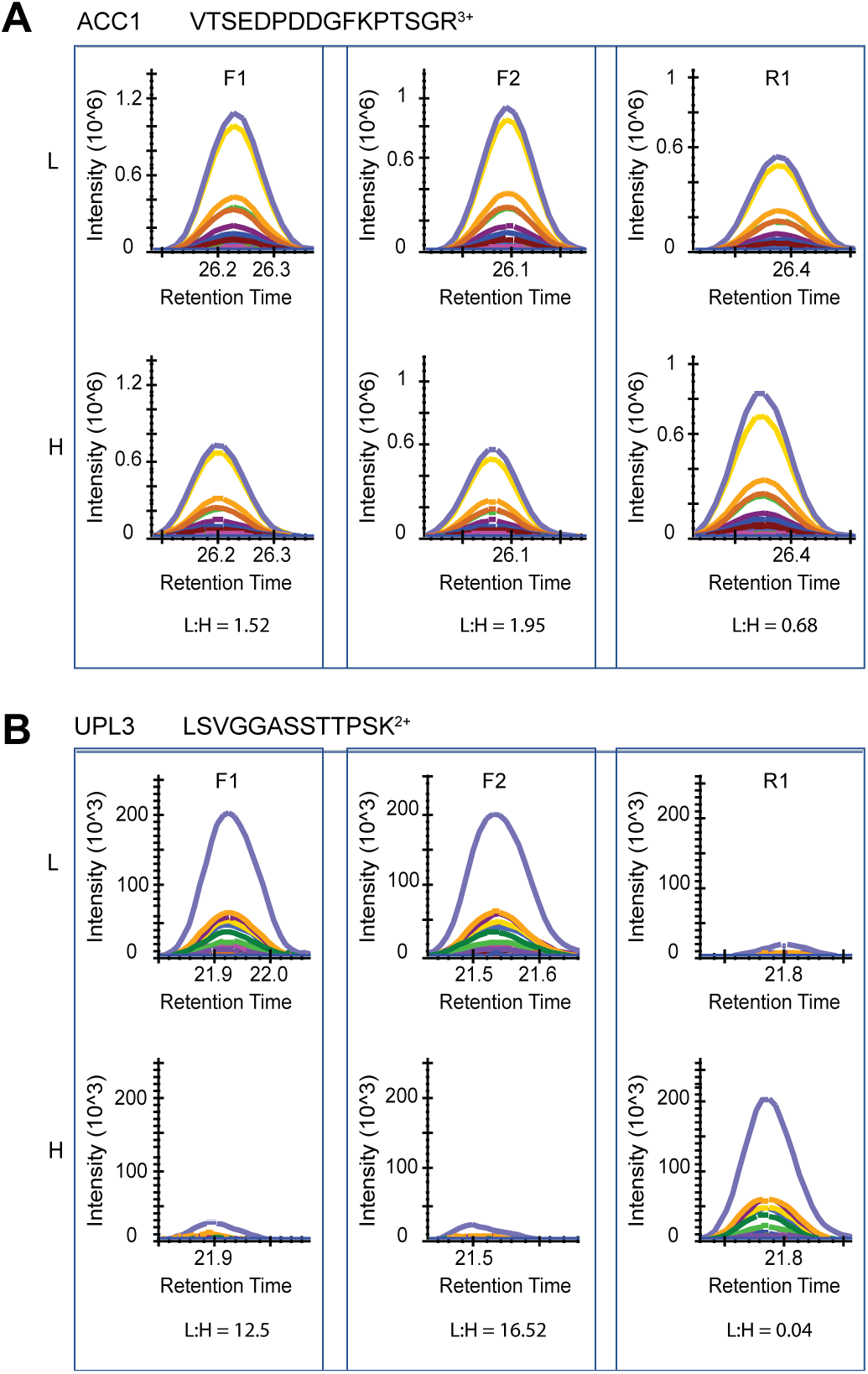
Targeted quantification of UPL3 and ACC1 in three BZR1 SILIP-MS experiments. PRM analysis of representative peptides of ACC1 (**A**) and UPL3 (**B**) in the light (^14^N, L) and heavy (^15^N, H) samples of the three replicates (F1, F2, and R1) of BZR1 SILIP-MS experiments. The *BZR1-YFP* sample was ^14^N-labeled (L) in forward replicates (F1 and F2), and ^15^N-labeled (H) in the reverse replicate (R1). Selected transitions were extracted from ^14^N and ^15^N labeled peptides of each replicate. The area under the curve of each transition was used for L/H ratio calculations.

**Figure S4.**
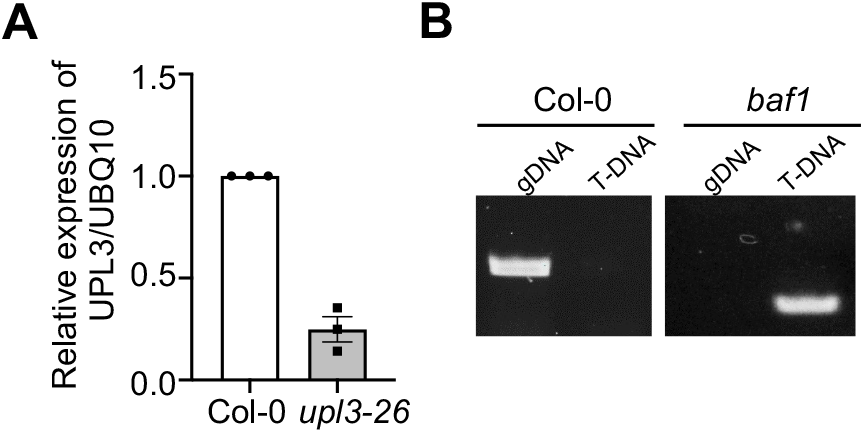
Characterization of the *upl3-26* and *baf1* mutants. **A)** The *UPL3* RNA level was analyzed by qRT-PCR, using *UBQ10* as the internal control. **B)** Genotyping of the *baf1* mutant.

